# Structural Host-Virus Interactome Profiling of Intact Infected Cells

**DOI:** 10.1101/2023.12.03.569778

**Authors:** Boris Bogdanow, Lars Mühlberg, Iris Gruska, Barbara Vetter, Julia Ruta, Arne Elofsson, Lüder Wiebusch, Fan Liu

## Abstract

Virus-host protein-protein interactions (PPIs) are fundamental to viral infections, yet high-resolution identification within the native context of intact infected cells has remained an unsolved challenge. Here, we developed structural host-virus interactome profiling (SHVIP) that combines *in situ* cross-linking mass spectrometry with the enrichment of newly synthesized viral proteins from infected cells. We established SHVIP using herpes simplex virus type 1 and obtained 739 PPIs based on 6,194 cross-links from productively infected cells. SHVIP captures PPIs across intracellular compartments and at the intact host endomembrane system. It resolves PPIs to the protein domain level and seamlessly integrates with AlphaFold-based structural modeling, facilitating detailed predictions of PPI sites within structured and intrinsically disordered regions. We show that SHVIP captures parts of the virus-host PPI space that are elusive to traditional interaction proteomics approaches. By selectively disrupting several newly identified virus-host PPIs, we confirm SHVIP’s ability to uncover genuine virus-host PPIs in the intact complex environment of infected cells.

## INTRODUCTION

Protein-protein interactions (PPIs) facilitate crucial processes during viral infections and are major determinants of viral pathogenesis. Identifying and characterizing these PPIs reveals potential drug targets and informs strategies to combat viral infections.

Various approaches allow screening for important host factors during viral infection. Insights into which host proteins enhance or impede viral replication have been gained through loss-of-function screens based on RNA interference ^1^ or CRISPR/Cas9 technologies ^2^. A possibility to capture physiologically relevant proteins at PPI level is affinity purification (AP) of tagged viral proteins and identification of the co-purifying interactors using mass spectrometry (MS) ^3, 4^. More recently, thermal proximity co-aggregation profiling (TPCA) ^5^ has been employed, which is based on the propensity of proteins within a complex to co-aggregate upon heat denaturation. For a variety of viruses, AP-MS ^6–16^ and TPCA ^17, 18^ have contributed functionally relevant insight into the virus-host interactome. However, these methods fail to capture PPIs from intact infected cells, cannot discriminate between direct and indirect interactors, and do not provide information on the interaction contact site ^4^. Furthermore, AP-MS requires non-native (i.e. tagged) protein baits typically introduced through transduction or transfection, making it labor-intensive and time-consuming to perform for many target proteins.

In principle, these limitations can be addressed by cross-linking mass spectrometry (XL-MS). XL-MS captures protein contacts using a small organic molecule (‘cross-linker’), typically composed of a spacer arm and two functional groups that are reactive towards specific amino acids such as lysines. A dedicated MS method and data analysis pipeline then identifies the interacting amino acid residues. These XL-MS data reveal PPIs and yield insights into *in situ* protein complex structures and binding interfaces. For example, they have successfully been used to validate ^19–21^ and augment ^22^ structural models obtained through AlphaFold2 (AF2) ^23^. Importantly, XL-MS is applicable to endogenous proteins in their native cellular environment and can provide PPI networks of intact organelles ^24, 25^, cells ^20, 26^, and viral particles ^27^. However, previous XL-MS studies on pathogen-host interactions from intact infected cells have been limited in sensitivity and identified fewer than 50 PPIs ^28^ likely due to the complexity and abundance of the background host proteome.

Virus infection is often accompanied by a pronounced downregulation of host protein synthesis, a phenomenon called “host shutoff” ^29^. Consequently, a large fraction of newly synthesized proteins in infected cells is of viral origin, which can be captured by metabolic pulse labeling with bio-orthogonal amino acids ^30–33^.

Here, we capitalize on this host shutoff mechanism by developing ***S****tructural **H**ost **V**irus **I**nteractome **P**rofiling (**SHVIP**),* which combines the bio-orthogonal labeling of newly synthesized proteins with proteome-wide XL-MS to sensitively map virus-host PPIs *in situ*. We applied SHVIP to herpes simplex virus type 1 (HSV-1), a medically relevant human herpesvirus with a complex proteome ^34^. The resulting HSV-1 interactome comprises hundreds of known and novel PPIs based on thousands of cross-links. It recapitulates PPIs within the intact subcellular context and resolves interactions at the sub-protein level. By integrating our SHVIP results with affinity purification, molecular genetics, and structural modeling, we deliver structural insight into dozens of host-virus PPIs.

## RESULTS

### SHVIP captures the viral interactome in intact HSV-1 infected cells

The SHVIP strategy presented here involves four steps **(Figure 1a)**. First, cells are infected with a virus that is able to replicate efficiently in a given cell line. Second, once cells are committed to a replicative viral life cycle, L-homopropargylglycine (HPG) is added to label newly synthesized - and thus predominantly viral - proteins for several hours. Third, the membrane-permeable cross-linker DSSO is added to the intact cells to covalently link proteins that are in proximity with each other. Fourth, HPG-containing proteins are extracted using copper-catalyzed click-chemistry together with their covalently linked interactors. Finally, the enriched proteins are digested, enriched for cross-linked peptides via strong cation exchange, and analyzed by MS.

**Figure 1.**
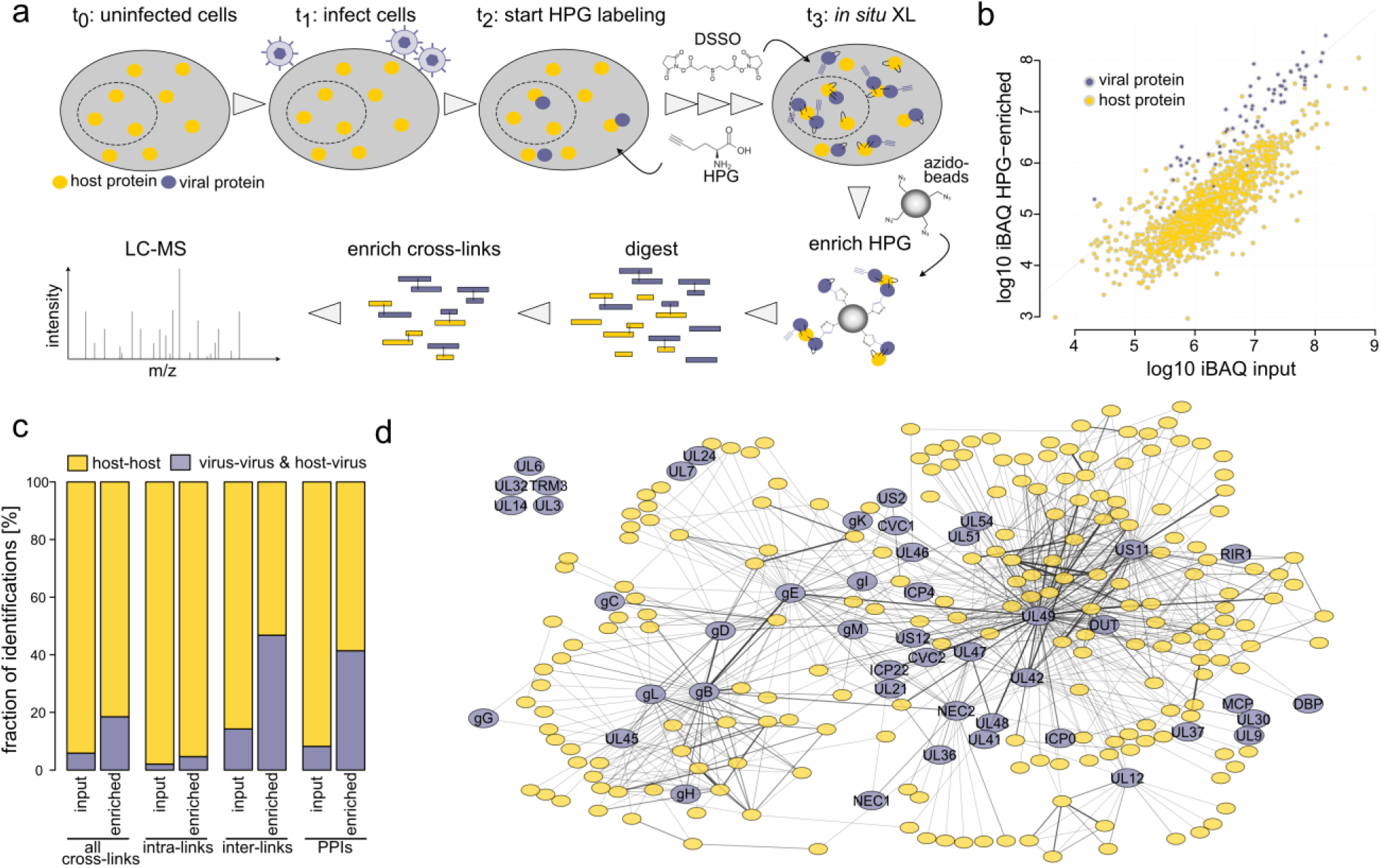
SHVIP workflow and proof of concept. **(a)** Workflow: Cultured cells are infected with a virus able to induce host shut-off. Once host shut-off is in place, newly synthesized proteins are labeled with L-HPG, and subsequently cross-linked with DSSO, and finally enriched using click chemistry. Proteins are digested into peptides, cross-links are enriched and measured by LC-MS. **(b)** Abundance (measured by iBAQ) of host and viral proteins in non-cross-linked HPG-enriched and input samples. **(c)** Relative fraction of identifications involving viral proteins or only host proteins for different identification types in XL-MS data. **(d)** Virus-centric PPI network of HSV-1 infected cells. Host proteins (yellow) are only included when they are directly linked to viral proteins (blue). Data are filtered to 1% separate cross-link FDR and reported at 1% inter-PPI FDR with 6,194 cross-links and 254 proteins. Edge width scales with the number of identified cross-links between the two proteins.

We applied this methodology to HSV-1 infected human embryonic lung fibroblasts (HELFs) and labeled the cells with HPG for 17 hours (7-24 hours post-infection). MS analysis of the linear (non-cross-linked) peptides in the enriched sample and an aliquot taken before HPG-enrichment (input) shows that HPG-enrichment reduces host protein abundance by a factor of 10 on average **(Figure 1b, Supplementary Table 1)**. Similarly, it increases the frequency of sequenced spectra of viral origin (**Extended Data Figure 1a**), and expands the fraction of viral proteins contributing to the overall intensity from ∼20% to ∼75% **(Extended Data Figure 1b)**. The contribution of viral proteins to the total protein intensity from cross-linked samples was slightly lower (∼60%) as additional cross-linked host proteins co-purified with the viral proteins. Importantly, SHVIP substantially increased the proportion of identified cross-links involving viral proteins **(Figure 1c)**. The highest increase was observed for cross-links between peptides originating from different protein sequences (inter-links) which is particularly valuable as inter-links give deeper insight into the viral PPI network. Both the relative frequency and absolute numbers of PPIs involving viral proteins increased upon HPG enrichment **(Extended Data Figure 1c)** and showed reproducibility similar to previous XL-MS studies from complex samples ^26^ **(Extended Data Figure 1d)**. Taken together, this demonstrates that SHVIP increases the sensitivity of viral interactome capture.

### Spatial reconstruction of host-virus PPIs

In order to provide a high-coverage structural interactome of the infected cell, we analyzed the combined enriched and input samples from two experiments involving different MS-acquisition schemes. We first filtered inter- and intra-links separately at a 1% residue pair-level false discovery rate (FDR). We then focused on the most relevant PPIs and generated a subnetwork of interactions among viral proteins and their direct host interaction partners (**Figure 1d**, **Supplementary Table 2, Extended Data Figure 2a**), yielding 739 interactions (of which 441 involve viral proteins) at a 1% separate PPI-level FDR estimated by target and decoy counts **(see Methods)**. These data cover 46 viral proteins, representing ∼65% of the viral proteins detected by MS-analysis of linear peptides. We observed cross-links for viral structural (e.g. tegument, glycoproteins) and non-structural proteins (e.g. replication machinery) as well as a few cross-links to the rigid nucleocapsid shell of the virus **(Extended Data Figure 2b**). Together, this indicates that SHVIP sensitively captures a large part of the available viral proteome.

To assess if SHVIP captures the virus interactome across the entire host cell, we performed unsupervised clustering based on centrality indices ^35^, dividing our network into six communities (**Figure 2a**). A GO-enrichment analysis, based on cellular component annotations of host proteins, revealed distinctly enriched categories for these communities (**Figure 2b**). While protein communities 4 and 5 enrich ribosomal proteins, community 6 contains many DNA binding proteins such as histones and other chromatin proteins (e.g. HMGB1, HMGB2, HMGN2). In two other communities (1 and 3), host proteins of the endomembrane system are enriched, such as proteins localizing to the endoplasmic reticulum (ER) lumen (e.g. B2M, CALR, P4HB, PDIA6, ERP44), ER-Golgi intermediate compartment or the Golgi apparatus (e.g. GALNT1, RAB6A, SCAMP2, SCAMP3). These data confirm that SHVIP can resolve host-virus PPIs across various cellular compartments.

**Figure 2.**
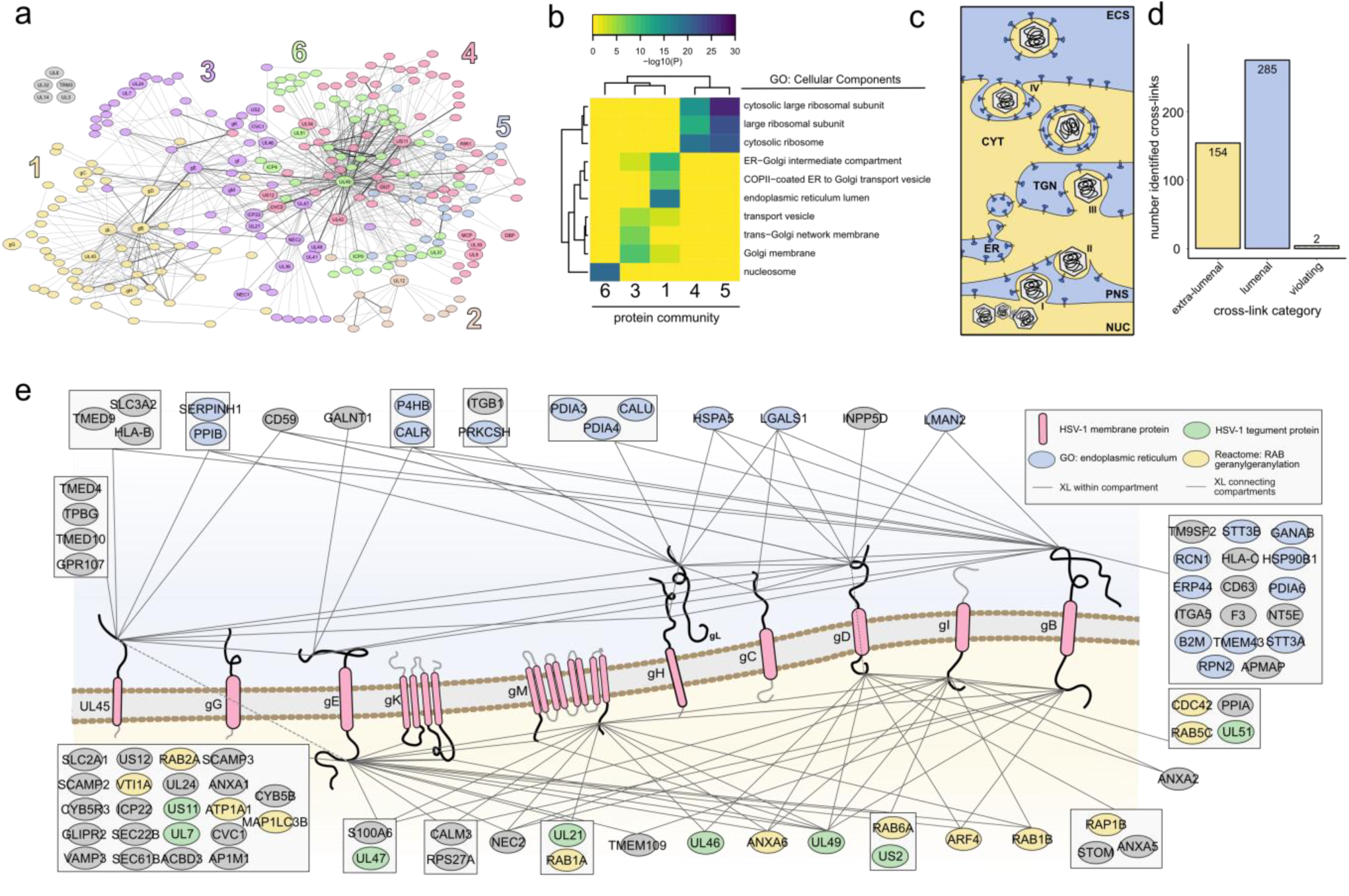
The structural interactome of HSV-1 at the host endomembrane system. **(a)** Unsupervised community clustering ^35^ of the structural interactome. Proteins colored according to the community numbers. **(b)** GO-enrichment based on cellular component annotations of host proteins within five of the communities. Community two had too few host proteins to enrich any GO term. **(c)** Extra-lumenal (yellow, e.g. cytosol) and lumenal (blue, e.g. ER lumen) compartments in late stage HSV-1 infection. **(d)** Number of cross-links connecting lumenal or extra-lumenal domains of viral transmembrane proteins and cross-linking partners with annotated domains. **(e)** Viral transmembrane proteins (red) shown in a membrane with their cytosolic tails pointing down and lumenal domains pointing up. Interaction partners with their domain-specific cross-links to either lumenal or extra-lumenal domains are grouped when they share cross-links to the same transmembrane protein domains. Color coding of host interaction partners according to GO annotations in legend. Host transmembrane proteins with several domain-specific links to viral transmembrane proteins are provided in Extended Data Figure 3a.

We were particularly interested in the PPIs at the endomembrane system. During the late stage of herpesvirus infection, cellular membranes become reorganized to allow assembly and release of mature virions. This is regulated by a complex network of PPIs, where membranes separate the involved proteins into a lumenal (of ER, trans-Golgi or vesicles) or extra-lumenal (cytosol, nucleoplasm or intra-virion) population (**Figure 2c**). While DSSO is able to pass the membrane, it is not able to cross-link proteins that are separated by membranes. Therefore, cross-links in this region allow evaluating whether SHVIP compromises the integrity of the cellular membranes. When evaluating cross-linking partners of viral envelope proteins at domain-level resolution **(Figure 2d)**, we found that almost all cross-links (99.6%) occurred between domains within either the lumenal or the extra-lumenal compartments, confirming that SHVIP captures interactions at a largely intact membrane system.

By mapping the domain-specific interaction partners onto HSV-1 transmembrane proteins **(Figure 2e, Extended Data Figure 3a)**, we found viral tegument proteins localizing to the extra-lumenal side of the membrane and interacting with cytosolic tails of viral glycoproteins, consistent with their ascribed role in mediating secondary envelopment ^36–38^. In addition, we observed many Rab GTPases, crucial regulators of endocytic and exocytic membrane trafficking, and related proteins at the cytosolic side. This includes not only RAB5 ^39^, RAB1A/B ^40^ and RAB6 ^41^ proteins with known proviral functions, but also RAB2A that is yet uncharacterized in HSV-1 infection **(Extended Data Figure 3b)**. At the lumenal side we observed many well-known ER-resident proteins, such as chaperones (PDIA4, PDIA6, HSP90B1) and glycosylation factors (GALNT1, GANAB), interacting with the extra-virion domains of viral glycoproteins, consistent with ER-luminal processing of viral glycoproteins. Interestingly, the viral membrane proteins gB and UL45 physically associated with cellular proteins involved in antigen presentation, such as HLA-B, HLA-C and B2M, potentially perturbing the cellular antigen presentation pathway^42^. In addition, glycoproteins gH/gL and gB, core components of the viral fusion machinery, were found cross-linked to alpha and beta subunits of integrins (ITGA5, ITGB1), which are known entry receptors for HSV-1 ^43^ **(Extended Data Figure 3c)**. We also observed cross-links between gH/gL and gB as well as the fusion-regulatory protein UL45 ^44^, supporting the idea of physical cross-talk among proteins of the HSV-1 fusion machinery^45^.

Similar to these viral glycoproteins, we also found other viral proteins that cause substantial membrane remodeling, such as the nuclear egress complex (NEC), composed of NEC1 and NEC2. NEC mediates the envelopment of nucleocapsids at the inner nuclear membrane, followed by their delivery to the outer nuclear membrane by membrane fusion ^46^. The *in situ* PPIs of this complex **(Extended Data Figure 3d)** reveal both known and novel insight into this process. First, we observed known functionally relevant interaction partners such as ICP22 ^47^ and EMD (also known as emerin) ^48^. Second, we found the extra-lumenal domains of several viral glycoproteins associated with NEC2, previously discussed as functionally important for fusion at the outer nuclear membrane ^49^. Third, we discovered the association of five ER-membrane proteins to the NEC, suggesting a yet unknown role of ER proteins during nuclear egress. Overall, SHVIP provides detailed, domain-level insights into host-virus interactomes at the endomembrane system, relevant for glycoprotein processing, membrane trafficking, adaptive immunity, egress and assembly.

### SHVIP confirms and extends AP-MS data

Having established that SHVIP provides extensive and biologically relevant data, we wanted to compare our approach to AP-MS, as the most widely employed method for virus-host PPI profiling ^6–16^. To this end, we selected eight viral proteins from our network, tagged them individually with an HA tag at their endogenous loci and performed anti-HA affinity purifications ^10^, enriching the bait with its interaction partners from infected cells. As controls, we used cells infected with the parental wildtype (WT) virus, not expressing any HA-tagged protein variants (**Figure 3a, Extended Data Figure 4a-h, Supplementary Table 3)**. On average, cross-linking partners of the bait show significantly higher enrichment in the corresponding anti-HA-bait precipitate than proteins with indirect or no cross-link connection to the bait (**Figure 3b,c**). At the level of individual baits, we observed significant differences in 6 out of 8 cases **(Extended Data Figure 4i-p)**. In total, at a stringent AP-MS log2 fold-change cut-off of 4, 42 SHVIP interactors could be validated by AP-MS **(Figure 3d)**. At a less stringent cut-off at log2 of 2, the number of overlapping interactors increased to 61 and remained significant **(Figure 3e)**. By utilizing a sliding cut-off, we observed a progressive decrease in the significance of the overlap when interactions below this threshold were considered **(Figure 3f)**. Thus, SHVIP reported PPIs can be validated by AP-MS and both techniques retrieve a common set of PPIs.

**Figure 3.**
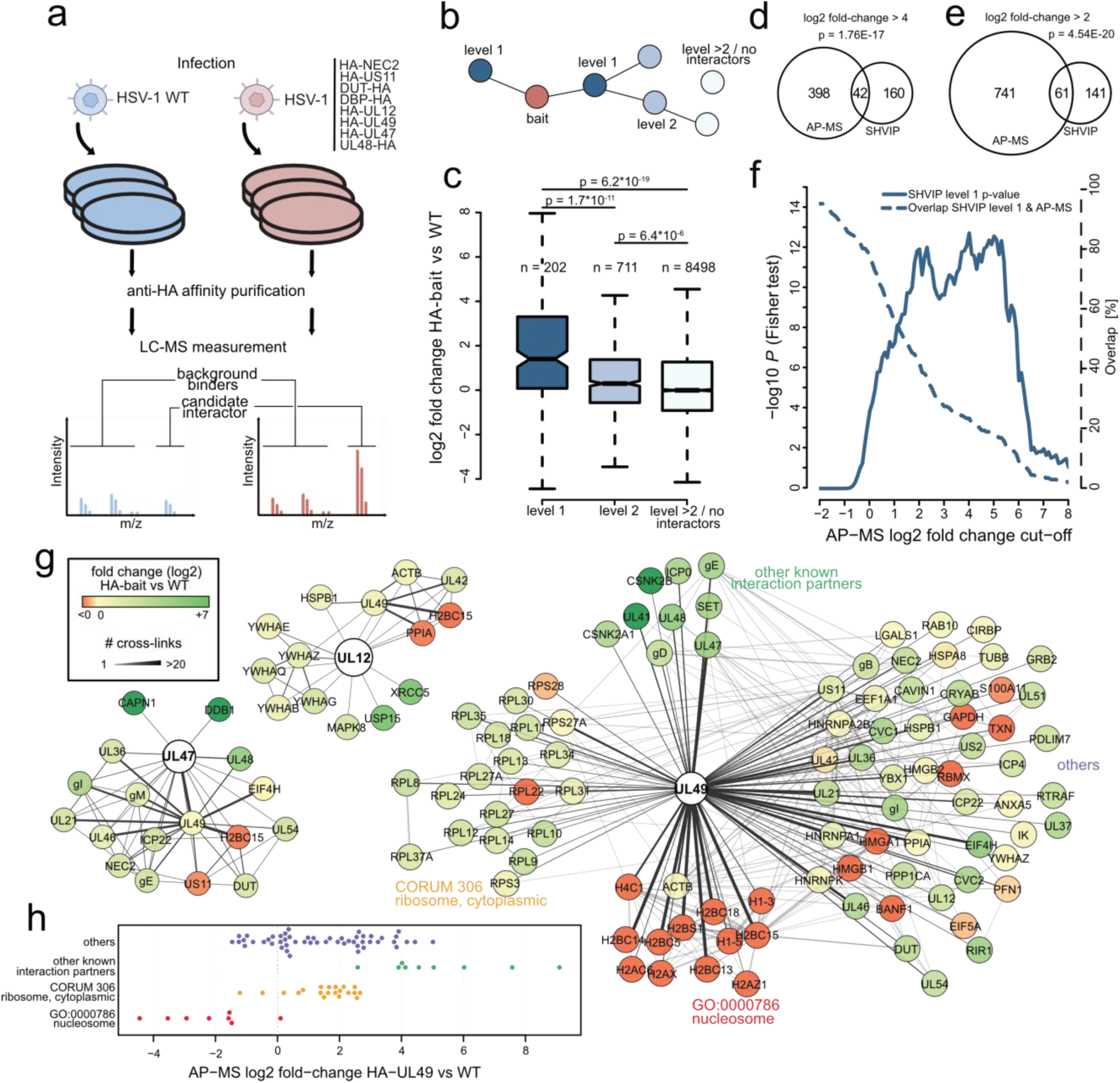
SHVIP complements and extends AP-MS data. **(a)** Workflow for AP-MS experiments. Transgenic HSV-1 viruses individually expressing the indicated HA-tagged protein variants were used to infect HELFs (n=3 biological replicates), followed by anti-HA enrichment and LC-MS measurement. Purifications were compared to control experiments with HSV-1 WT not expressing any HA-tagged bait. The intensity ratio comparing HA-bait to control enrichments was computed for individual co-enriched proteins (Volcano plots for all baits are included in Extended Data Figure 4). **(b)** For comparing to AP-MS data, proteins in the SHVIP interactome (see also Figure 1d) were categorized into proteins cross-linked to the bait (level 1), proteins cross-linked to the bait via one other protein (level 2), or higher-order interactors and proteins not contained in the structural interactome (>level 2/ no interactors). **(c)** Log2 fold-changes in the respective AP-MS experiment comparing the protein categories from panel b. P-values are based on a two-sided wilcoxon rank-sum test. **(d,e)** Comparison of the interactome of the 8 selected viral proteins in SHVIP and AP-MS at a higher-stringency (d) and lower-stringency (e) log2 fold-change cut-off applied to AP-MS data. An additional t-test P-value cut-off of 0.01 was applied to potential AP-MS interactors. The displayed P-values indicate the statistical significance of the overlap between proteins identified via AP-MS and SHVIP and are based on one-sided Fisher’s Exact test. **(f)** Comparing the overlap between SHVIP and AP-MS and its statistical significance using a sliding AP-MS fold-change cut-off. P-values are based on one-sided Fisher’s Exact test. **(g)** Representative depiction of SHVIP interactomes for selected proteins. Color coding of interacting nodes according to the co-enrichment ratios with the bait in AP-MS. **(h)** AP-MS co-enrichment levels of UL49 interactors found by SHVIP. Interactors are grouped into different biological categories.

Looking at individual protein examples shows that SHVIP and AP-MS capture overlapping as well as complementary parts of the interactome. For the viral proteins UL12 and UL47, almost all cross-linking partners were also found enriched in anti-HA precipitates **(Figure 3g)**. In the case of UL49, an abundant tegument protein with a wide variety of functions and interaction partners ^50^, we observed a more mixed picture when comparing AP-MS and SHVIP: Several known UL49 interaction partners, such as casein kinase 2 ^51^, UL41 (also known as vhs), UL48 (also known as VP16) ^52^ and SET (also known as TAF-I) ^53^ were detected by XL-MS and were highly enriched in AP-MS **(Figure 3g, h)**. However, other biologically relevant UL49 interactors detected by XL-MS, e.g. ribosomal proteins (consistent with UL49’s known polysome association ^54^), displayed only intermediate AP-MS co-enrichment levels. Moreover, nucleosome-associated proteins were observed as the most frequent cross-linking partners of UL49, corroborating microscopy data ^55–57^ and *in vitro* experiments ^56, 57^. Surprisingly, they were not enriched in AP-MS, suggesting that the nucleosome-UL49 interaction is disrupted by cell lysis or the enrichment procedure. An explanation could be that these interactions are linked to UL49 undergoing liquid-liquid phase separation with DNA^58^, a concentration-dependent process that is disturbed by cell lysis. These results suggest that, while many PPIs are identified by both SHVIP and AP-MS, those depending on the *in situ* environment of the infected cell are uniquely captured by SHVIP.

### Integrating SHVIP with AF2 delivers system-wide structural insights

In addition to identifying interacting proteins in the infected cell, SHVIP can provide structural insights into the cross-linked protein assemblies. This is possible as cross-linkers bridge a defined distance range, e.g. up to 35 Å for the DSSO cross-linker applied here. As such, our SHVIP data can support the structural characterization of heterodimers from HSV-1 infected cells.

Only three inter-links (2 PPIs) could be mapped onto existing experimental structures of heterodimers **(Figure 4a)**, reflecting the paucity of structural information on HSV-1-host interactions. In order to extend insights into yet unsolved heterodimeric structures, we predicted all binary interactions in our dataset by AF2-Multimer^23^ **(Figure 4b)** and categorized model quality based on the calculated docking scores (pDockQ)^59^. In general, dimers between host proteins yielded predictions with better scores than those between viral proteins only or between viral and host proteins **(Figure 4c)**. This is consistent with the comparably low success rates of docking predictions for viral proteins ^59^, lower depth in the multiple sequence alignment (MSA) for viral proteins and lower expected co-evolutionary signals captured in MSA between two organisms (host and virus). In addition, when comparing AF2-models to cross-links, we noticed that inter-links involving viral proteins occurred more frequently in poorly predicted (pLDDT < 50), likely intrinsically disordered regions (IDRs) than those between host proteins **(Figure 4d)**.

**Figure 4.**
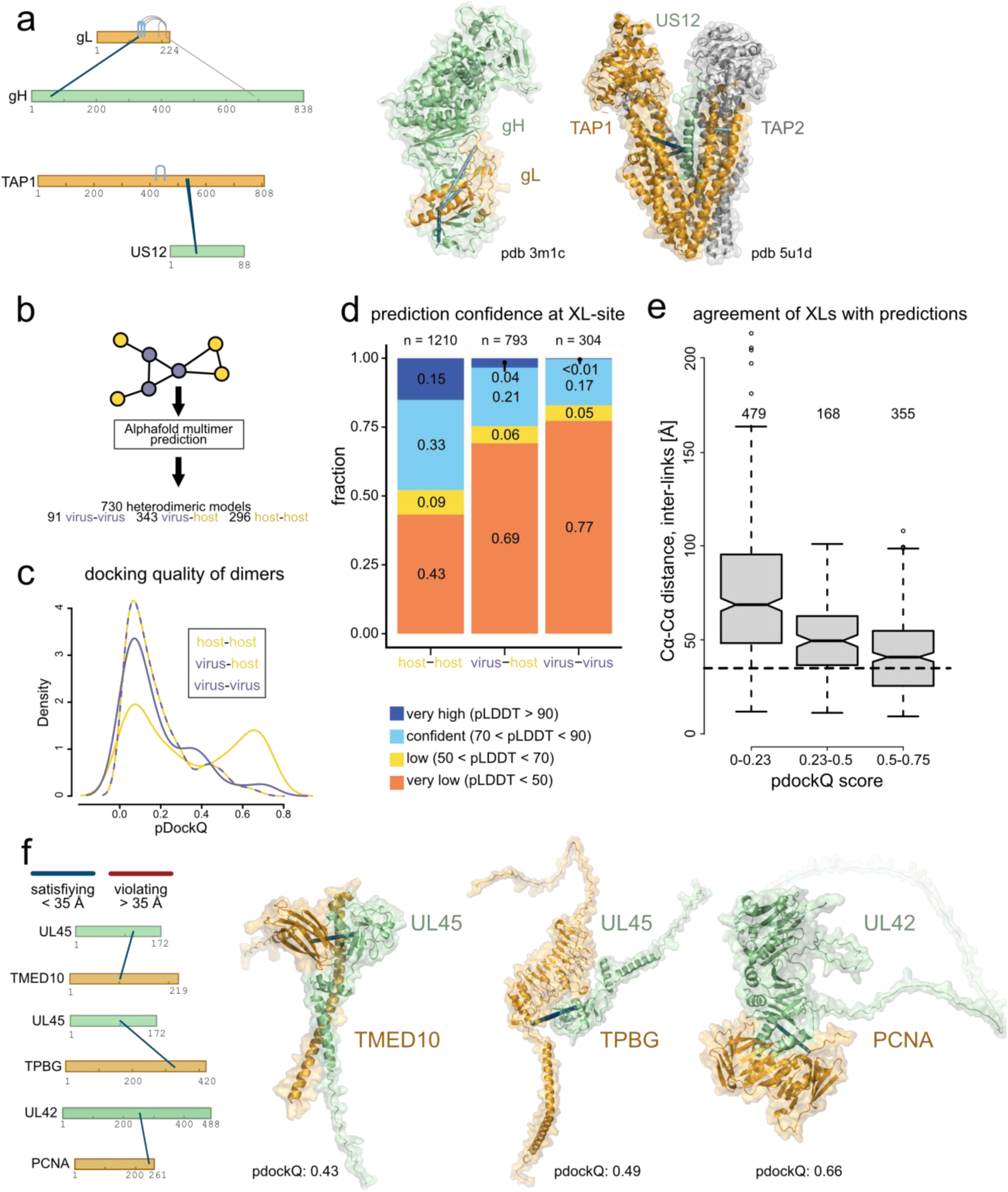
Structural insight into host-virus PPIs through SHVIP and AF2. **(a)** Cross-links mapped onto heterodimers with experimentally solved structures. Proteins shown as bars ^89^ or high-resolution experimental structures (gH-gL, pdb 3m1c, and US12-TAP1/2, pdb 5u1d). Cross-links highlighted gray could not be mapped to the experimental structures because of missing coordinates. All other cross-links meet the DSSO distance constraint of 35 Å. **(b,c)** Predicting virus-virus, host-virus and host-host dimers of the structural interactome using AF2-Multimer **(b)** with their respective pDockQ distributions **(c). (d)** Prediction confidence (AF2-Multimer pLDDT score) at the specific inter-linked lysine residue for different types of inter-links. The lower pLDDT on either lysine residue of the cross-link was used for categorization. **(e)** Distribution of inter-link distances (Cα-Cα distance of cross-linked lysines) for AF2-Multimer models in different pDockQ score ranges. Shown are only inter-links involving lysines from protein regions with pLDDT scores > 50. **(f)** Examples of well-predicted models agreeing with the cross-linker distance constraint.

We initially focused on structural models with inter-links in well-predicted regions (pLDDT > 50). In these cases, cross-links provide orthogonal experimental evidence to validate AF2-Multimer predictions. Indeed, the distances bridged by the cross-linker decreased with increasing pDockQ **(Figure 4e)**, indicating that the better the model quality, the better the cross-link distances fit into the predicted structures.

In sum, we obtained 68 models (18 involving viral proteins) that have both acceptable docking scores (pDockQ > 0.23) and an agreement of at least half of the inter-links with the prediction **(Supplementary Table 4, Figure 4f)**. Among them, we modeled the viral transmembrane protein UL45 with transmembrane and lumenal domain contacts to the ER-resident transmembrane protein TMED10, which regulates the secretion of interleukin 1-beta and other leaderless cargoes ^60^. UL45 was also adequately docked to the cell surface domain of another transmembrane protein, the Wnt1/β-catenin signaling inhibitor TPBG^61^. Further, SHVIP data supported the structural prediction for UL42, the viral polymerase processivity subunit, binding to its cellular homolog PCNA ^62^. This interaction may explain why PCNA is enriched at viral DNA replication forks ^63^.

We more specifically investigated the dimer between the tegument protein UL47 and DDB1 **(Figure 5a)**. DDB1 functions as an integral component of Cullin 4-RING ubiquitin ligase (CRL4) complexes ^64^. Two UL47-DDB1 cross-links were identified, with one of them satisfying the distance constraint in the corresponding AF2 model. In the model, the UL47 C-terminal region in proximity to the satisfied inter-link adopts an alpha-helical fold, in positional congruence with previously described viral DDB1 interactors, such as RCMV-E27 **(Figure 5b)** ^65^. To confirm the UL47-DDB1 interaction and assess the contribution of the UL47 C-terminal helix, we performed co-immunoprecipitation (IP) experiments. While DDB1 efficiently co-precipitated with UL47-WT, deleting the C-terminal 18 amino acids (676-93) of UL47 was sufficient to disturb this interaction **(Figure 5c)**. In contrast, UL47-UL48 binding ^66^ was not affected. Further AP-MS experiments confirmed that the UL47 C-terminal truncation leads to a strong and specific loss of DDB1-CUL4A binding **(Figure 5d)**, indicating that the C-terminal helix acts as a critical DDB1 interaction module.

**Figure 5.**
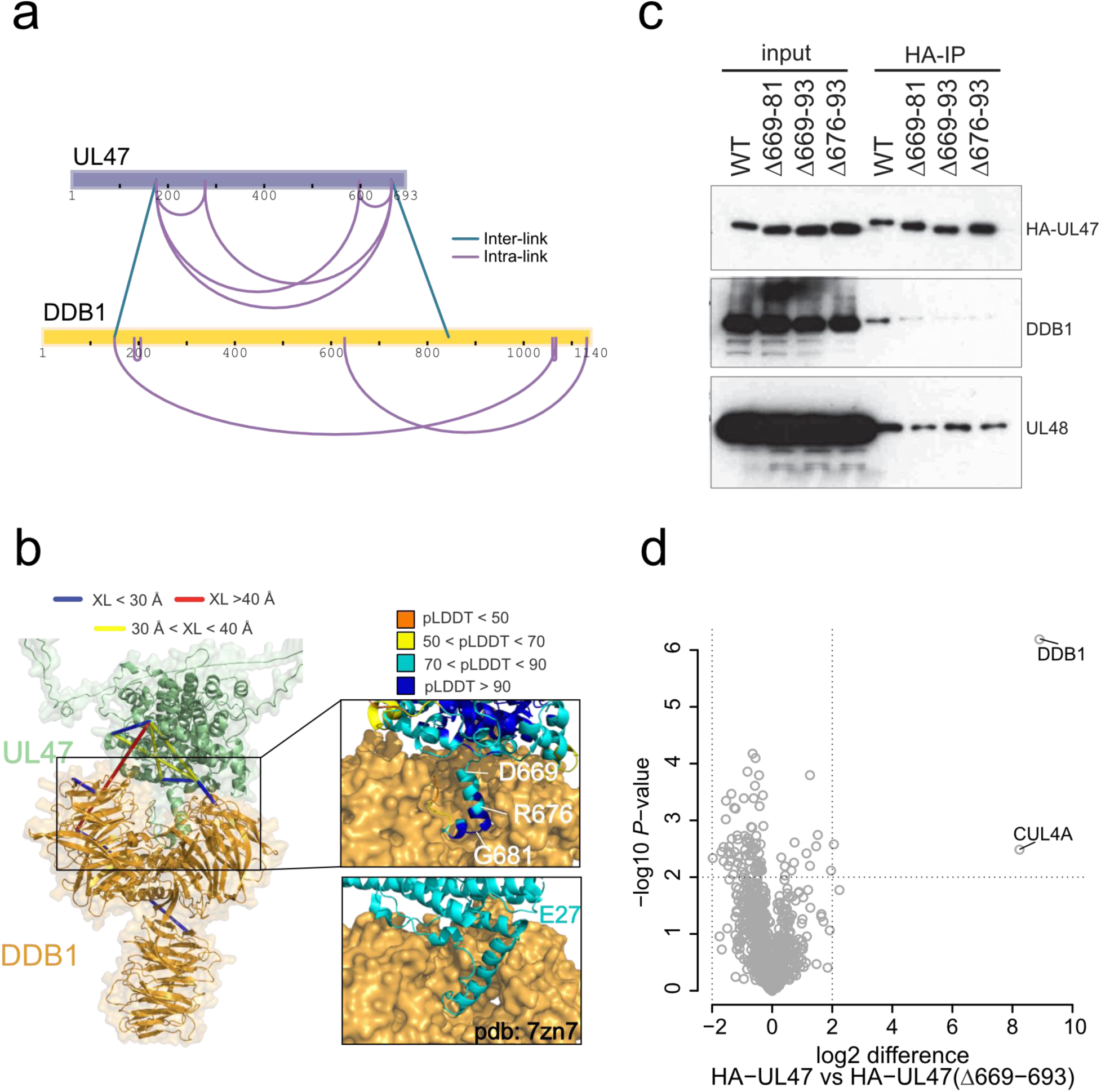
Structural determinants of the DDB1-UL47 heterodimer. **(a)** UL47-DDB1 cross-links mapped onto sequence bars ^89^. **(b)** DDB1-UL47 cross-links mapped onto the predicted dimer with color coding of the cross-links according to distances between Cα atoms. The inset shows the AF2 prediction confidence for the C-terminal helix of UL47. Displayed below the inset is the experimentally resolved structure of E27-DDB1 (pdb 7zn7) for comparison. **(c)** HA-directed Co-IPs against different mutant and WT variants of UL47 from infected cells at 24hpi. **(d)** AP-MS experiments directly comparing the interactome of WT UL47 to mutant UL47 in a label-free set-up based on n=3 replicates.

Thus, combining SHVIP with AF2 enables the prediction of a compendium of virus-host/ virus-virus dimers, which can be validated by cross-linking distance constraints, reverse genetics, and orthogonal interaction assays.

### Identifying sequence determinants for virus-host PPIs in IDRs

Most of our inter-links occurred in poorly predicted regions, likely IDRs. These regions are hotspots for short linear motifs (SLiMs) that viruses use to interact with host proteins^67^. To find such interaction sites, we screened all AF2-multimer models of our cross-linked virus-host PPIs for short stretches of ordered amino acids (pLDDT > 50) that are in contact with the respective binding partner and in a disordered regional context **(Figure 6a)**. We obtained a list of 28 candidates **(Supplementary Table 5)** with varying average confidence scores and amino acid length **(Figure 6b)**. We manually curated this list to yield 12 sites, representing a promising set of putative SLiMs mediating host-virus interactions.

**Figure 6.**
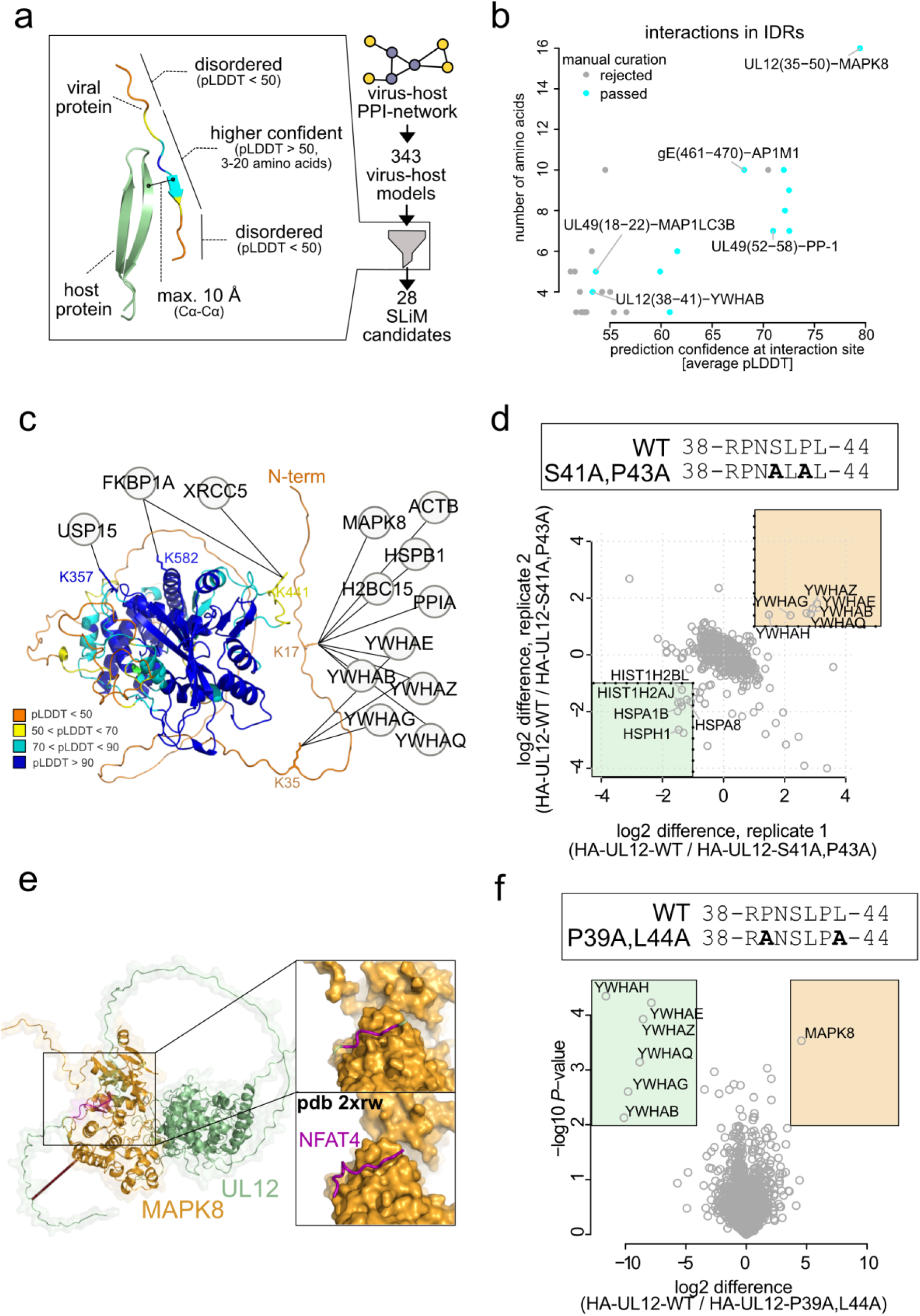
Identifying sequence determinants for host-virus interactions in IDRs. **(a)** Workflow for selecting putative interaction sites in IDRs from AF2 models. **(b)** Comparison of the residue length for the putative interaction motif to the confidence of the predicted interface (average pLDDT of all amino acids within the selected site). **(c)** Host proteins cross-linked to UL12, depicted on the monomeric AF2 model of UL12, color coded by pLDDT. **(d)** Mutational disruption of the MAPK8 binding site in UL12 in the viral genome of HSV-1, creating S41A, P43A-UL12. SILAC-based comparative AP-MS of the WT UL12 to the S41A, P43A mutated UL12 interactome, with both (n=2, label-swap) replicates depicted. Boxed regions highlight proteins that interact stronger with S41A, P43A-UL12 (shaded green) or WT-UL12 (shaded orange), based on a log2 fold-change cut-off of 2 in both replicates. **(e)** Predicted MAPK8-UL12 dimer with a putative D-motif highlighted magenta. The inset shows the magnified region of the D-motif in UL12 interacting with the D-site in MAPK8. Shown below the inset is a solved crystal structure of MAPK8 with the D-motif containing peptide of NFAT4 (pdb 2xrw). **(f)** Mutational disruption of the D-motif in UL12 in the viral genome of HSV-1, creating P39A, L44A-UL12. Label-free comparative AP-MS of the WT UL12 and P39A, L44A-U12 interactomes. The p-values are based on two-sided t-tests without multiple hypothesis correction of n=3 replicates. Boxed regions highlight proteins that interact significantly stronger with P39A, L44A-UL12 (shaded green) or WT-UL12 (shaded orange), based on a cut-off at P=0.01 and log2 fold-change of 4.

The best predicted site was observed in a short stretch of amino acids in the N-terminal disordered domain of the alkaline nuclease UL12 (amino acids 35-50) binding to MAPK8 kinase (also known as JNK1). Curiously, we observed another predicted interaction site to the 14-3-3 protein YWHAB within the same region of UL12 (amino acids 38-41). Both MAPK8 and 14-3-3 proteins were also cross-linked to N-terminal lysines within UL12 (K35, K17) **(Figure 6c)**.

We hypothesized that both MAPK8 and 14-3-3 may bind to the same short sequence stretch within UL12 and thus more closely evaluated the predictions, starting with YWHAB-UL12. The AF2 model puts amino acids 38 to 41 of UL12 into the peptide binding channel of the 14-3-3 protein **(Extended Data Figure 5a)**. This is consistent with S41 being suggested as the best hit for 14-3-3 interaction using a sequence-based prediction tool ^68^ **(Extended Data Figure 5b)**. Following these two lines of evidence, we mutated critical amino acids in the viral genome (S41A, P43A) to disrupt 14-3-3 binding **(Extended Data Figure 5c, Figure 6d)**. We then compared WT to mutant in an AP-MS experiment using SILAC (stable isotope labeling of amino acids in cell culture) labeling. WT UL12 co-precipitated with 14-3-3 proteins stronger than S41A, P43A-UL12. In contrast, the mutant interactome showed enrichment of several heat-shock proteins and histones, suggesting effects on UL12 folding or stability upon loss of 14-3-3 binding.

In the MAPK8-UL12 dimer model, the disordered N-terminal domain of UL12 wraps around the MAPK8 kinase domain and binding occurs between a D-motif in the UL12 unstructured region and the MAPK8 docking groove **(Figure 6e)** ^69^. We mutated the candidate D-motif (P39A, L44A, **Figure 6f)**, leaving the 14-3-3 motif intact. Comparing mutant to WT UL12 interactomes indicates that the mutations result in loss of MAPK8 binding but stronger association with other UL12 interactors (14-3-3 proteins). Thus, our analysis identifies two short linear motifs that co-exist on a narrow stretch of amino acid within the UL12 N-terminal disordered domain and enable UL12 to recruit MAPK8 and 14-3-3 proteins.

In addition, our list of putative SLiMs includes other compelling predictions. For example, we found the interaction of gE with the adaptor protein AP1M1, with a putative tyrosine-trafficking motif in gE **(Extended Data Figure 6a)**. Also, we predicted the central player in autophagy MAP1LC3B (also known as LC3-B) interacting with UL49, which has a putative LC3 interaction region (LIR) ^70^ **(Extended Data Figure 6b)**. Further, we observed the interaction between UL49 and the cellular Phosphatase PP-1 (also known as PPP1CA), for which the predicted model indicated binding of an amino acid stretch in the unstructured region of UL49 to the RVXF-motif recruitment site in PP-1 **(Extended Data Figure 6c)**. As the corresponding amino acid sequence in UL49 suggests the presence of a non-canonical RVXF motif **(Extended Data Figure 6d)**, we aimed for independent validation of this binding site. Indeed, we found that mutating the motif in UL49 (R52A, F57A) reduces PP-1-UL49 binding in HA-directed AP-MS assays **(Extended Data Figure 6e,f)**. These findings showcase that combining SHVIP and AF2-based structure predictions allows the discovery of critical sequence determinants of virus-host PPIs within intrinsically disordered regions.

## DISCUSSION

While interaction proteomics has contributed immensely to uncover the molecular processes underlying viral pathogenesis ^3, 4, 6–18^, a global view of interaction contact sites within infected cells has remained elusive. The SHVIP approach introduced here addresses this shortcoming. Using SHVIP, we define the structural interactome of a complex herpesvirus in intact infected cells, revealing distinct clusters of spatially separated virus-host PPIs. Orthogonal validations, structure predictions, and molecular genetics provide insights into the structural basis of a subset of these interactions.

SHVIP integrates well with AF2 to inform structural predictions at interactome scale ^19–21^ delivering residue-residue connections as corroborative evidence to purely *in silico* analyses. Enhancing interactome-wide AF2 modeling with SHVIP is particularly valuable from a virological point of view because (1) AF2-only predictions of dimers involving viral proteins have generally lower quality than those of host proteins ^59^ **(Figure 4)** and (2) experimental structures of host-virus protein complexes are scarce for HSV-1 and many other viruses. Augmenting AF2 with SHVIP strengthened 18 protein complex predictions, which are potentially linked to DNA replication, immune evasion, and signaling. For instance, we found a C-terminal helix in the tegument protein UL47 that interacts with the CRL4 adapter unit DDB1 **(Figure 5)** in a manner that is typical for DCAF-type substrate receptors of CRL4. It will be interesting to see whether UL47 acts in analogy to other viral DCAF mimics ^65^ and exploits CRL4 to target antiviral host proteins for proteasomal degradation.

SHVIP also allows characterizing viral proteins that are challenging to study outside their cellular context. This includes proteins undergoing liquid-liquid phase separation ^71^ (e.g. UL49), transmembrane proteins ^72^ (e.g. viral glycoproteins), or proteins involved in membrane remodeling, like the NEC. SHVIP can capture weak and context-sensitive PPIs because it capitalizes on protein cross-linking within intact cells. In contrast, such interactions may not be amenable to AP-MS or TPCA ^17, 18^ as they involve cell lysis. Furthermore, AP-MS is primarily based by protein affinity whereas XL-MS is driven by protein proximity. These methodological differences likely explain why both techniques provide overlapping information while also offering a substantial degree of complementarity **(Figure 3)**.

We observed that cross-links involving viral proteins are enriched in IDRs **(Figure 4d)**, recognized hotspots for SLiMs frequently exploited by viruses to manipulate host processes ^67^. Based on SHVIP-supported heterodimers and their corresponding AF2-models, we propose 12 potential SLiMs within IDRs **(Figure 6a).** We validated the non-canonical RVXF-motif in the UL49 protein as a SLiM for binding PP-1 phosphatase. In analogy to HCMV-UL32 ^27^, UL49-PP-1 interaction may be important to regulate the tegument phosphorylation state^73^, facilitating virion assembly. Furthermore, we identified two co-existing SLiMs within the disordered segment of alkaline nuclease UL12 responsible for competitive binding of MAPK8 and 14-3-3 proteins. These interactions might constitute a regulatory switch, allowing HSV-1 to link the viral replication machinery to MAPK8, a potent effector of cellular stress signaling that is crucial for reactivating HSV-1 from latency ^74^. These examples show that canonical and non-canonical interaction motifs are discoverable by integrating SHVIP and AF2. The distinctive advantage of this approach lies in its incorporation of structural context and co-evolutionary relationships, enabling a significant improvement over sequence-based methods for predicting SLiMs^68, 75^.

SHVIP is, in principle, applicable to study native cells infected with any virus inducing host shutoff ^29^. This includes medically or veterinary relevant viruses such as influenza A viruses ^30^, poxviruses ^76^, picornaviruses ^77^, flaviviruses ^78^, or asfarviruses ^33^. Also, it does not require upfront genetic work, which makes it readily applicable to emerging viruses and viruses that do not tolerate epitope tagging in their genome. However, it is important to consider that HPG-labeling requires methionine starvation, impacting cell viability^79^ and viral titers^80^, particularly when employed for prolonged periods. This may in the future be mitigated by adopting starvation-free protocols ^79^ for SHVIP.

In summary, SHVIP offers various conceptual advantages for virus-host PPI profiling. Our data on a complex and medically relevant herpesvirus deliver a blueprint for the structural characterization of PPIs within their intact environment during viral infections. The method unlocks an opportunity for the in-depth characterization of viral gene functions during cellular infections.

## MATERIAL & METHODS

### Cells and Viruses

Human embryonic lung fibroblasts (HELFs) were maintained as previously described ^81^ and used for preparation of viral stocks. Recombinant HSV-1 strain 17 ^82^ was used for all experiments. Infectious virus titres were determined by immunotitration and indicated as immediate early (IE)-forming units (IU) per mL. In brief, HELFs were infected with serial dilutions of cell-free virus stocks. At 8 hpi, the number of infected cells was determined by flow cytometry of IE antigen ICP4 expression. Viral mutants were created by traceless bacterial artificial chromosome mutagenesis according to established protocols ^83^. Mutations were verified by Sanger sequencing and PCR. See Supplementary Table 6 for a list of mutagenesis primers.

### Stable isotope labeling in cell culture

HELFs were cultured for at least five passages in SILAC-DMEM medium supplemented with 10% (v/v) dialysed fetal bovine serum (FBS, Pan-Biotech), 1 x Glutamax (Gibco), 1 x non-essential amino acids (Gibco), 100 Units/mL Penicillin (Gibco) and 100 µg/mL Streptomycin (Gibco). For “heavy” labeling medium was supplemented with 0.8 mmol/L L-[13C6,15N2]-Lysine (Lys8) and 0.4 mmol/L L-[13C6,15N4]-Arginine (Arg10). For “light” labeling medium was supplemented with natural Lysine and Arginine to the same concentrations respectively.

### Co-immunoprecipitation

Cells were harvested at 24 h post infection and extracted by freezing-thawing in immunoprecipitation buffer, containing 50 mM Tris–Cl pH 7.4, 150 mM NaCl, 10 mM MgCl_2_, 1 mM NaF, 10 mM β-glycerophosphate, 0.5 mM Na_3_VO_4_, 10 mM N-ethylmaleimide, 0.5% Nonidet P-40 (NP-40), 10% glycerol, 1 mM dithiothreitol (DTT), 0.3 μM aprotinin, 23 μM leupeptin, 1 μM pepstatin, 0.1 mM Pefabloc. HA-UL47-containing protein complexes were immunoprecipitated by incubating extracts with HA-specific antibodies and protein G-coupled sepharose beads (4 Fast Flow, GE Healthcare). The sepharose-bound proteins were analyzed by standard immunoblotting for the presence of HA-UL47, DDB1 and UL48. The following antibodies were used: anti-DDB1 (rabbit polyclonal, Bethyl Laboratories, A300-462A), anti-HA (rat clone 3F10, Roche, ROAHAHA), anti-UL48 (mouse clone 1-21, Santa Cruz Biotechnology, sc-7545).

### Affinity-purification mass spectrometry

Eight viral baits were selected from our network for subsequent orthogonal validation by AP-MS. We excluded viral glycoproteins, capsid or capsid-associated proteins and proteins with fewer than two PPIs. From the remaining shortlist we selected three viral proteins with few PPIs (<10 PPIs: UL48, DBP, DUT), three with moderate number of PPIs (>10 and <50 PPIs: UL12, NEC2, UL47) and two highly inter-linked proteins (UL49, US11). AP-MS was carried out essentially as described previously ^10^. Label-free experiments were performed in triplicates and SILAC-based experiments were performed in label-swap duplicates. In all cases cells were harvested 24 h after infection with WT or mutant virus. One fully confluent 15 cm dish of HELFs per replicate and condition was used. Following washing with PBS, cells were lysed by a 30 min incubation in lysis buffer: 25 mM Tris-HCl (pH 7.4), 125 mM NaCl, 1 mM MgCl_2_, 1% NP-40, 0.1% sodium dodecyl sulfate (SDS), 5% glycerol, 1 mM DTT, 0.3 μM aprotinin, 23 μM leupeptin, 1 µM pepstatin, 0.1 mM Pefabloc. Lysates were cleared by centrifugation and the cleared lysates were incubated with anti-HA magnetic microbeads (Miltenyi) for 60 min before being applied to µMACS microcolumns (Miltenyi). We used lysis buffer for the first washing step, lysis buffer without detergent for the second and 25 mM Tris-HCl (pH 7.4) for the final washing step. Protein material was eluted in a total volume of 0.2 mL 8 M guanidine hydrochloride at 95 °C and precipitated from the eluates by adding 1.8 mL LiChrosolv ethanol and 1 µL GlycoBlue. After incubation at 4 °C overnight, samples were centrifuged for 1 h at 4 °C and ethanol was decanted before samples were subjected to sample preparation.

### SHVIP sample preparation

Cells were infected at a multiplicity of infection of 5 IU/cell, using standard protocols. At 7 hours post infection, cells were washed twice with pre-warmed PBS before the L-HPG-labeling medium was added, consisting of 500 µM HPG (Click Chemistry Tools), methionine-free DMEM (Gibco ref.) and 10% dialyzed fetal bovine serum (Pan Biotech). Cells were washed in PBS and harvested by scraping at 24 hpi before being cross-linked with 5 mM DSSO in a 1:1 (vol/vol) PBS/cell suspension for 1 h at room-temperature under constant shaking. Afterwards, cells were lysed in a buffer containing 200 mM Tris (pH 8.0), 4% 3-((3-cholamidopropyl) dimethylammonio)-1-propanesulfonate, 1 M NaCl, 8 M urea and protease inhibitors. Cell lysates were incubated for 30 min at 4°C. Genomic DNA in the samples was digested by addition of Benzonase. Afterwards, cell lysates were sonicated in a Bioruptor Pico (Diagenode) with 10 cycles of 30 s sonication pulses, followed by 30 s without sonication. Samples were centrifuged at 10,000 g for 10 min and the resulting pellet was discarded. Approximately 5% of the sample volume was saved as input while the remainder was subjected to enrichment of labeled proteins.

HPG-labeled proteins were enriched using copper(I)-catalyzed azide-alkyne cycloaddition also known as “click reaction”, using the Click-&-Go protein enrichment kit (Click Chemistry Tools) according to the manufacturer’s protocol. In brief, samples were incubated over-night rotating together with picolyl-azide conjugated agarose beads and a copper(I)-ion containing catalyst solution. Protein-conjugated agarose beads were centrifuged at 1,000 g for 1 min and the supernatants containing unbound, unlabeled cellular proteins were discarded. Afterwards, disulfide protein bridges were reduced by adding 10 mM DTT and alkylated by adding 40 mM chloroacetamide (CAA). Subsequently, agarose beads were transferred to 0.8 mL Pierce centrifuge columns (Thermo Scientific). Beads were subjected to a five-step washing protocol employing 10 times 500 µl of the following washing buffers: i) 1% w/v SDS, 250 mM NaCl, 5 mM EDTA in 100 mM Tris (pH 8.0); ii) 8 M urea in 100 mM Tris (pH 8.0); iii) 80% v/v acetonitrile (ACN) in water; iv) 5% v/v ACN in 50 mM triethylammonium bicarbonate (TEAB); v) 2 M urea plus 5% v/v ACN in 50 mM TEAB. After the last washing step, beads were resuspended in the final washing buffer. To release the captured proteins from the beads, proteins were digested by addition of trypsin and lysyl endopeptidase C (Lys-C). Samples were incubated at 37°C overnight shaking. The proteolytic digestion was stopped by addition of 1% formic acid (FA). The peptide-containing supernatants were recovered by 5 min centrifugation at 500 g. Peptides were desalted using Sep-Pak C8 1cc vacuum cartridges (Waters) according to manufacturer’s protocol. Proteins from input samples were extracted using methanol-chloroform precipitation. Precipitated proteins were resuspended in a buffer containing 1% w/v SDC, 5 mM Tris-(2-carboxyethyl)phosphin (TCEP), 40 mM CAA in 50 mM TEAB (pH 8.0). The following digestion by trypsin and Lys-C was conducted as described above.

### Off-line fractionation

Desalted peptides were further fractionated by strong cation exchange chromatography (SCX) using an Agilent 1260 Infinity II HPLC system equipped with a PolySULFOETHYL A column (PolyLC). Peptides were separated using a 95 min gradient ranging from 100% Buffer A (20% ACN in water) to a mixture of 20% Buffer A and 80% Buffer B (5 M NaCl in 20% ACN and water), collecting 45 s fractions. Fractions were desalted using C8 stage tips ^84^ and dried by speed vacuum. Dried samples were stored at −20°C before LC-MS measurement.

### AlphaFold2 predictions

Hetero-dimers were predicted by AlphaFold2 multimer v2.1 ^85^ using the default protocol and database sequence search methods. The DDB1-UL47 hetero-dimer was additionally predicted using AF2.3 parameters and used for display in Figure 4. The AF2 model of HSV1-UL12 was downloaded from https://www.bosse-lab.org/herpesfolds/.

The pDockQ scores were calculated with previously published parameters ^59^ and model quality was considered based on the pDockQ score (0-0.23: poor, 0.23-0.5: acceptable, 0.5-0.75: good). Mapping of cross-links onto AlphaFold2-Multimer predictions was performed using the bio3d R package. Therefore, we extracted the C-alpha atom coordinates of inter-linked lysines in the best predicted dimeric structure and calculated their distance in three-dimensional space. For matching models to cross-link data, we first filtered for models with a pDockQ score higher than 0.23. We then re-ranked AF2 models of the same dimer by prioritizing in order: (i) highest number of cross-links from structured regions (pLDDT >50), (ii) the highest number of satisfied cross-links and (iii) the highest pDockQ score. Models with at least half of cross-links in-range (< 35 Å) were included in Supplementary Table 4.

To identify interaction sites within disordered regions, we developed a two-step algorithm in Python 3.10 utilizing Biopython ^86^. For each predicted dimeric structure (all ranks), residues of a chain are extracted if their corresponding C-alpha coordinates exhibit a Euclidean distance of less than 10 A to C-alpha coordinates of the other chain. Following this, we investigated these stretches of residues for their pLDDT scores. As a pLDDT below 50 is a predictor of disorder ^87^, consecutive residues with a motif length of three to up to 20 amino acids are extracted if each of them exhibits a pLDDT above 50 and if they are adjacent to residues with a pLDDT below 50.

### LC-MS of XL-MS data

Cross-linked peptides were measured on an Orbitrap Fusion Lumos Tribrid system (Thermo Fisher Scientific) equipped with a FAIMS Pro Duo interface (Thermo Fisher Scientific) operating with Xcalibur 4.6 and Tune 4.0. For this, approximately 1 µg of sample was loaded with an online-connected Ultimate 3000 RSLC nano LC system (Thermo Fisher Scientific) onto a 50 cm analytical, in-house packed reverse-phase column (Poroshell 120 EC-C18, 2.7 µm, Agilent Technologies) and separated with a 180 min or 120 min gradient going from 0.1% w/v FA in water (buffer A) to 0.1% w/v FA in 80% v/v ACN (buffer B) at a flow rate of 250 nl/min. FAIMS compensation voltages were alternated between -50, -60 and -75 V.

For the SHVIP experiment with MS2-only acquisition scheme, cross-linked peptides were detected using a stepped-HCD-MS2 method, where for MS1 Orbitrap resolution was set to 120,000 with a scan range of 375-1600 m/z and mass range set to “Normal”. Standard parameters were used for automatic gain control (AGC) and 50 ms of injection time was used. Precursors of charge states 4-8 were selected with an isolation window of 1.6 m/z and fragmented by stepped-HCD with the energies 21%, 27% and 33%. Dynamic exclusion was enabled with an exclusion period of 60 s. Cycle time in between master scans was set to 2 s. For MS2, Orbitrap resolution was set to 60,000 with mass range set to “Normal” and scan range to “Auto”. Maximum of injection time was adjusted to 118 ms and AGC target to 200%.

For the SHVIP experiment with the MS2-MS3 acquisition scheme, cross-linked peptides were detected by a stepped HCD-MS2-CID-MS3 method. MS1 and MS2 scan parameters were the same as for the stepped-HCD-MS2-only method. MS3 scans were triggered when DSSO signature peaks with a mass difference of 31.9721 mass units were detected. The 2 most intense reporter peaks with charge states 2-6 were selected as precursors for MS3 with an isolation window of 2 m/z and fragmented by CID with a collision energy of 35% and an activation time of 10 ms. MS3 scans were acquired in the Ion Trap with scan rate set to “Rapid”, 100 ms injection time, AGC target set to 200%, mass range set to “Normal” and scan range set to “Auto”.

### LC-MS of bottom-up data

Bottom-up proteomic samples were measured on an Orbitrap Fusion Tribrid instrument (Fusion), an Orbitrap Fusion Lumos instrument (Lumos), an Orbitrap Elite instrument (Elite) (all Thermo Fisher Scientific) connected online to an Ultimate 3000 RSLC nano LC system (Thermo Fisher Scientific) or an Orbitrap Exploris 480 (Exploris) connected online to a Vanquish neo UHPLC system (Thermo Fisher Scientific). Peptides were separated on an in-house packed 50 cm analytical, reverse-phase column (Poroshell 120 EC-C18, 2.7 µm, Agilent Technologies) with 120 min or 180 min gradients at a flow rate of 250 nl/min.

Methods run on Fusion or Lumos were measured with a HCD-MS2 method with MS1 scans acquired in the Orbitrap with the resolution set to 120,000, while MS2 spectra were acquired in the Ion Trap. Cycle time in between master scans was set to 1 s and dynamic exclusion was set to 40 s. Intensity threshold was set to 1E04 for Fusion and to 5E03 for Lumos and maximum injection time to 50 ms, while AGC target was kept at standard settings. Precursors of charge state 2-4 were selected for fragmentation with a precursor isolation window of 1.6 m/z and fragmented with the HCD energy of 30%.

For measurement on Exploris, peptides were measured with a HCD-MS2 method with MS1 scans acquired in the Orbitrap with the resolution set to 120,000. Cycle time in between master scans was set to 2 s and dynamic exclusion was set to 40 s. Intensity threshold was set to 1E04 and maximum injection time was set to “Auto”. AGC target was set to 300%. Precursors of charge state 2-4 were selected for fragmentation with a precursor isolation window of 1.6 m/z and fragmented with the HCD energy of 30%. MS2 spectra were acquired in the Orbitrap with a resolution set to 15,000 with AGC target set to “Standard” and maximum injection time set to “Auto”.

Measurements run on Elite were acquired using a CID-MS2 method. MS1 scans were acquired in the Orbitrap with resolution set to 120,000 and mass range from 350 to 1500 m/z. Dynamic exclusion was enabled with a duration of 60 s. Top 15 precursors of charge state 2 and 3 were selected with an isolation window of 1 m/z for fragmentation by CID. Normalized collision energy of 35%, activation time of 10 ms and activation Q of 0.25 were used for CID fragmentation. MS2 spectra were acquired in the Ion trap with mass range set to “Normal” and scan rate set to “Rapid”.

### Cross-link data analysis

Peak lists (.mgf files) were generated in Proteome Discoverer (v.2.1) to convert .raw files into .mgf files containing HCD-MS2 or HCD-MS2-MS3 data, respectively. The .mgf files were searched using a stand-alone search engine based on XlinkX v.2.0 ^88^ with the following settings: MS ion mass tolerance, 10 ppm; MS2 ion mass tolerance, 20 ppm; fixed modification, Cys carbamidomethylation; variable modification, Met oxidation; enzymatic digestion, trypsin; allowed number of missed cleavages, 3; DSSO cross-linker, 158.0038 Da (short arm, 54.0106 Da; long arm, 85.9824 Da). Spectra were searched against a concatenated target-decoy database generated on the basis of the host-virus proteome determined by bottom-up proteomics. Raw files from both experiments (MS2-only and MS2-MS3 acquisition) were searched separately, then combined and the FDR at 1% was applied at the level of residue-residue connections, separately for intra- and inter-links on the basis of a target-decoy competition strategy using randomized decoys. Following this, all cross-links were retained when they matched to a viral protein or to host protein with a viral interaction partner. The PPI-level FDR in this set of PPIs was estimated based on the ratio of decoys to targets. The resulting network was clustered with the edge-betweenness algorithm ^35^ using in-house R scripts and the edge.betweenness.community function from igraph package. Clusters and networks were visualized in Cytoscape v3.7.2 or using xiNET ^89^. The comparison of PPI and cross-link identifications between input and enriched samples are based on individual searches as described above, but with a naive 1 % FDR (that is, FDR filtering applied on the combined set of intra- and inter-links).

### Bottom-up proteomics data analysis

AP-MS experiments were designed as triplicate experiments comparing anti-HA APs in lysates containing the transgene to control purifications of the WT strain, or in case of mutation of short linear interaction motifs comparing HA-tagged WT protein containing virus against the HA-tagged mutant protein containing virus. The set-up was label free and control experiments were performed and measured in parallel. Raw-files were analyzed using MaxQuant v.2.0.3.0 ^90^, with standard parameters and match between runs, iBAQ and LFQ enabled. The HSV-1 reference proteome UP00009294 and Uniprot reference proteome of human protein sequences (downloaded 2020) was used. FDR cut-offs were set at 1% at PSM, protein, and modification site level. The proteinGroups.txt file was used for subsequent analysis with potential contaminants, reverse database hits and proteins only identified by a modification site removed. LFQ intensities were log2 transformed and, when appropriate, missing values were imputed on the basis of a normal distribution shrinked by a factor of 0.3 and downshifted by 1.8 standard deviations only when a protein was quantified in all three replicates of either experiment or control. Log2 fold-changes and p-values from a t-test were calculated based on these values. The significance of overlap between AP-MS and XL-MS was calculated using phyper function in R.

SILAC-quantified AP-MS experiments were conducted in duplicates comparing HA-tagged WT target protein containing virus with HA-tagged mutant target protein containing virus. Raw-files were analyzed using MaxQuant v. 1.6.2.6. Quantification was based on MaxQuant normalized SILAC-ratios using two parameter groups in the analysis for light and heavy (Lys8 and Arg10) labeled proteins, respectively. Remaining parameters were kept as described above, and the re-quantify option was enabled. The proteinGroups.txt file was used for data evaluation in R with log2 transformed SILAC ratios. Proteins enriched or depleted over a log2 ratio of 1 in both replicates were considered as hits.

## DATA AVAILABILITY

The mass spectrometry proteomics data have been deposited to the ProteomeXchange Consortium via the PRIDE ^91^ partner repository with the dataset identifier PXD047422 (reviewer details Username, Password: XXXXXX).

## AUTHOR CONTRIBUTIONS

B.B. conceptualized the project. B.B., L.M., I.G., J.R., L.W. and F.L. developed the methodology. B.B., L.M., I.G., B.V., J.R., A.E. and L.W. conducted the investigations. B.B., L.M., I.G. and L.W. conducted formal analysis. I.G., B.V., L.W., A.E. and F.L. procured resources. B.B. and L.M. performed visualization. B.B. and L.M. curated the data. B.B., L.W. and F.L. acquired funding. B.B., L.W. and F.L. administered the project. B.B., L.W. and F.L. supervised the project. B.B. and L.M. wrote the original draft. B.B., L.W. and F.L. reviewed and edited the manuscript.

## ACKNOWLEDGMENTS

The authors thank Beate Sodeik (MHH, Hannover) for providing us with recombinant HSV-1 strain 17 and Philip Lössl (Absea Biotechnology, Berlin) for critically reviewing and editing the manuscript. Funding was provided by the Deutsche Forschungsgemeinschaft (DFG) grant BO 5917/1-1 (B.B.). L. M and F. L. were supported by Leibniz-Wettbewerb (P70/2018). Financial support to A.E. was obtained from the Swedish Research Council for Natural Science, grant No. VR-2016-06301 and Swedish E-science Research Centre and from Knut and Alice Wallenberg Foundation. Computational resources to A.E. was obtained by the Swedish National Infrastructure for Computing via grants: SNIC 2021/5-297, SNIC 2021/6-197, Berzelius-2021-29 and Berzelius-2022-106.

## EXTENDED DATA FIGURES AND LEGENDS

**Extended Data Figure 1.**
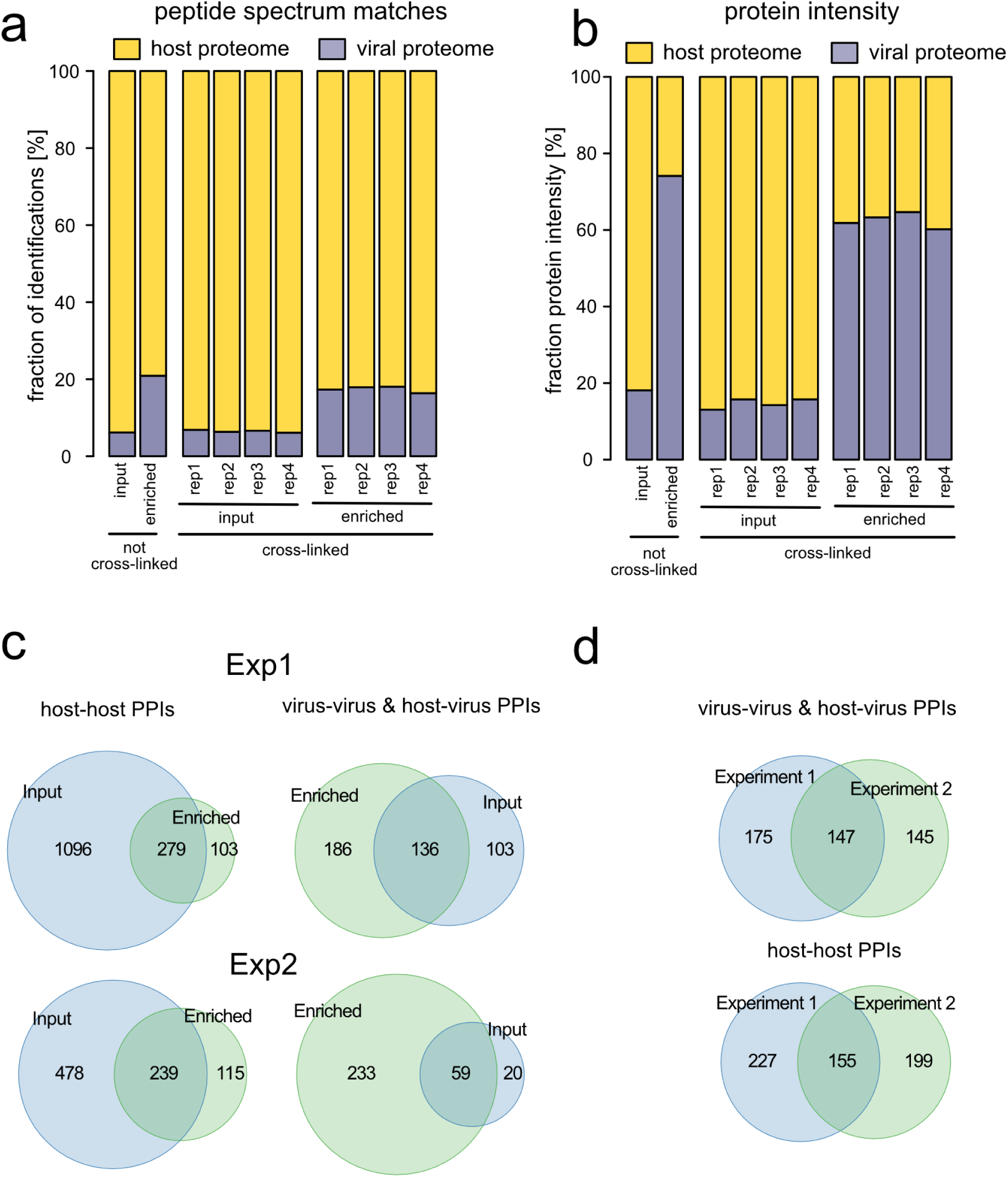
SHVIP quality control. **(a,b)** The number of peptide spectrum matches (a) and overall protein intensity (b) from an analysis of the linear peptides by LC-MS/MS with indicated treatments. **(c)** Overlap of PPI identifications in enriched and input samples after cross-link analysis in both experiments. Experiment 1 was performed in MS2-MS3 and experiment 2 in an MS2 only acquisition scheme. **(d)** Venn diagrams showing reproducibility of PPI identifications between HPG-enriched samples from both experiments.

**Extended Data Figure 2.**
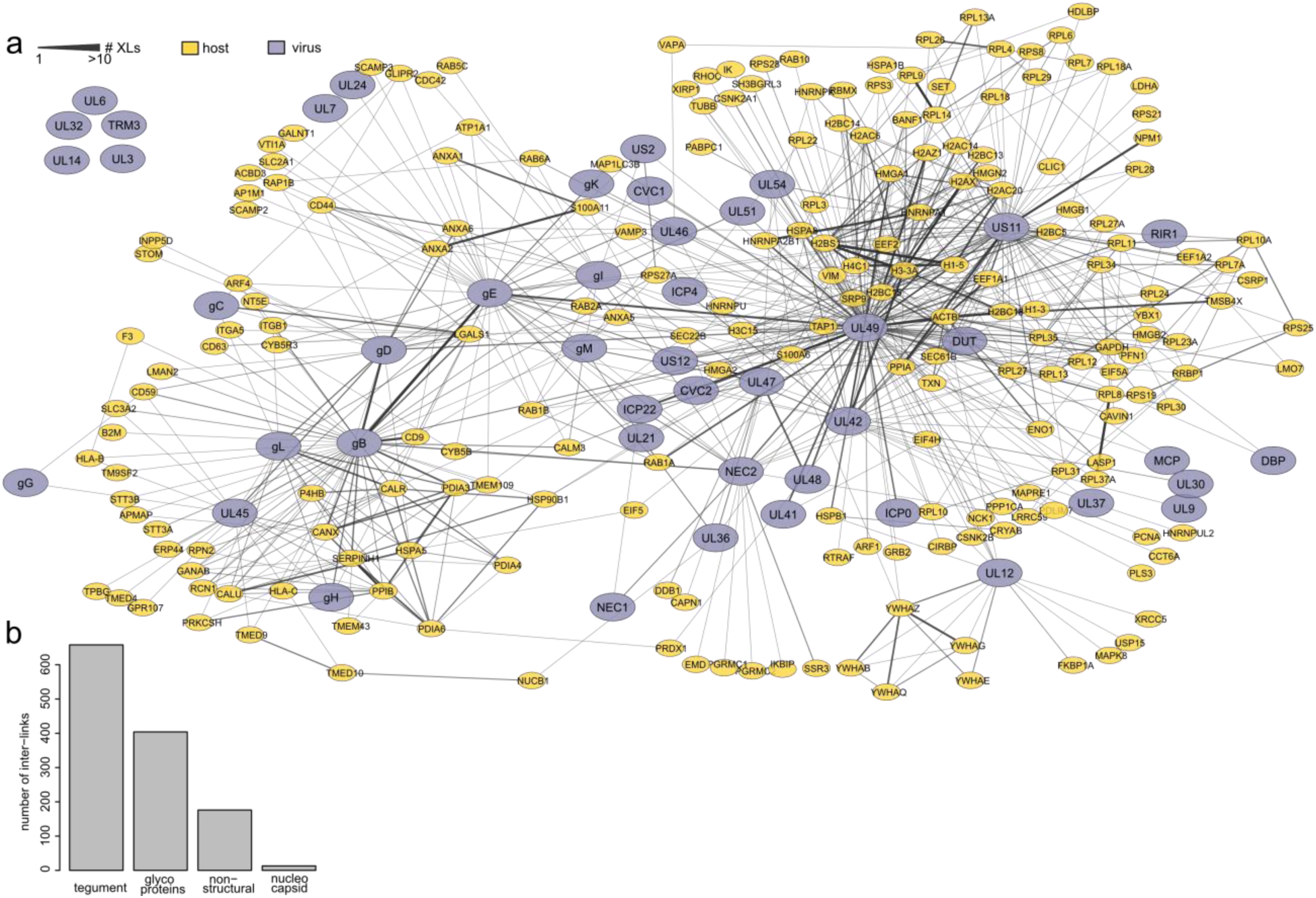
The structural interactome of HSV-1 infected intact cells. **(a)** Network of the structural interactome of HSV-1 infected cells with the Gene names for host and viral proteins (see also Figure 1d). **(b)** Inter-link coverage for viral proteins from different classes.

**Extended Data Figure 3.**
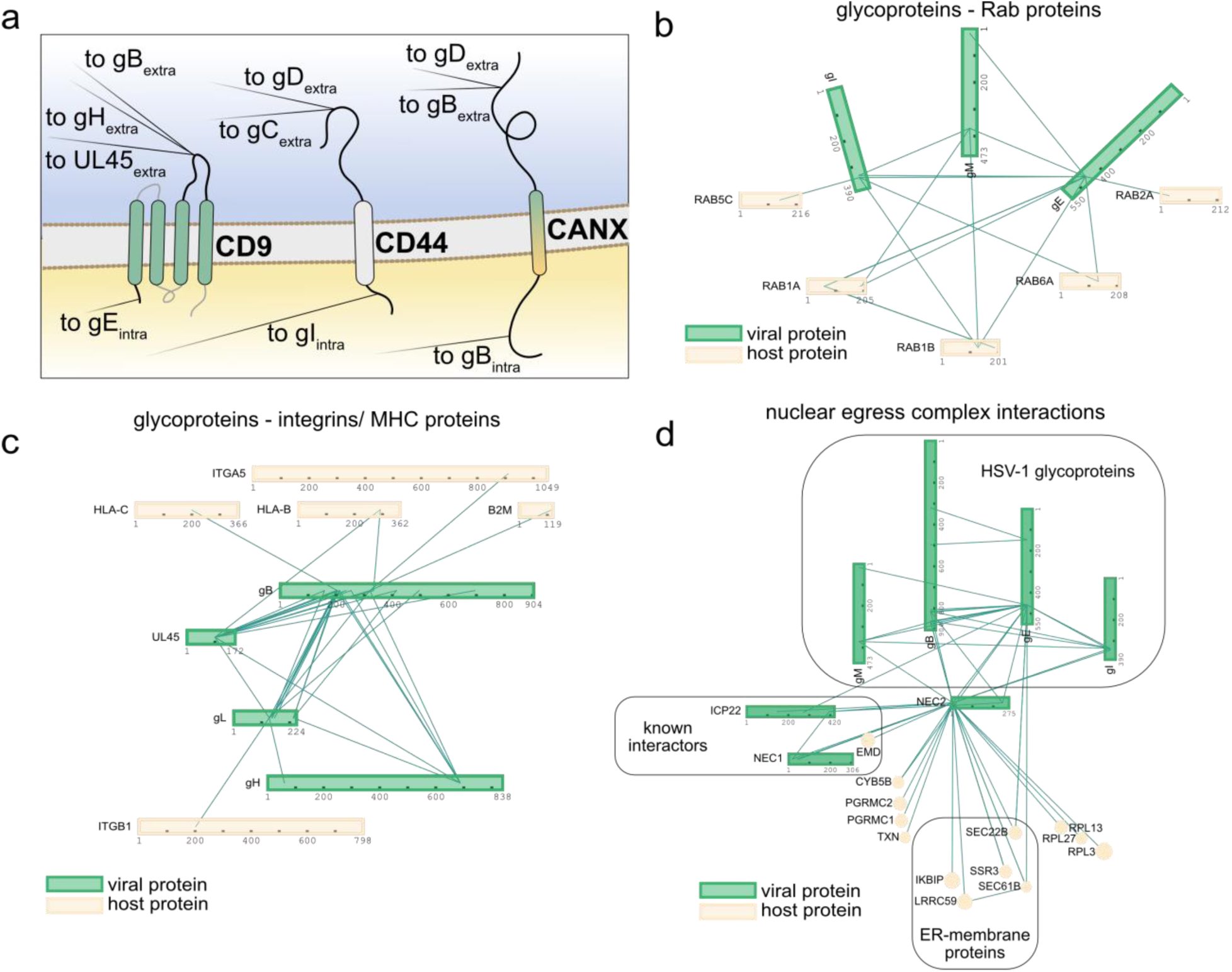
Selected Subnetworks at the host membrane system. **(a)** Three transmembrane host proteins cross-linked to viral transmembrane proteins via their lumenal and extra-lumenal domains. Black edges indicate cross-links to viral transmembrane proteins (Extra: lumenal domain, intra: extra-lumenal domain). **(b-d)** Selected subnetworks ^89^ of RAB proteins and their viral transmembrane protein interaction partners **(b)**, integrins, MHC proteins and their viral transmembrane protein interaction partners **(c),** and NEC2 with cellular and viral interaction partners **(d)**. The NEC2 interactors UL49, UL47 and US11 are not included in panel d for reasons of visual clarity.

**Extended Data Figure 4.**
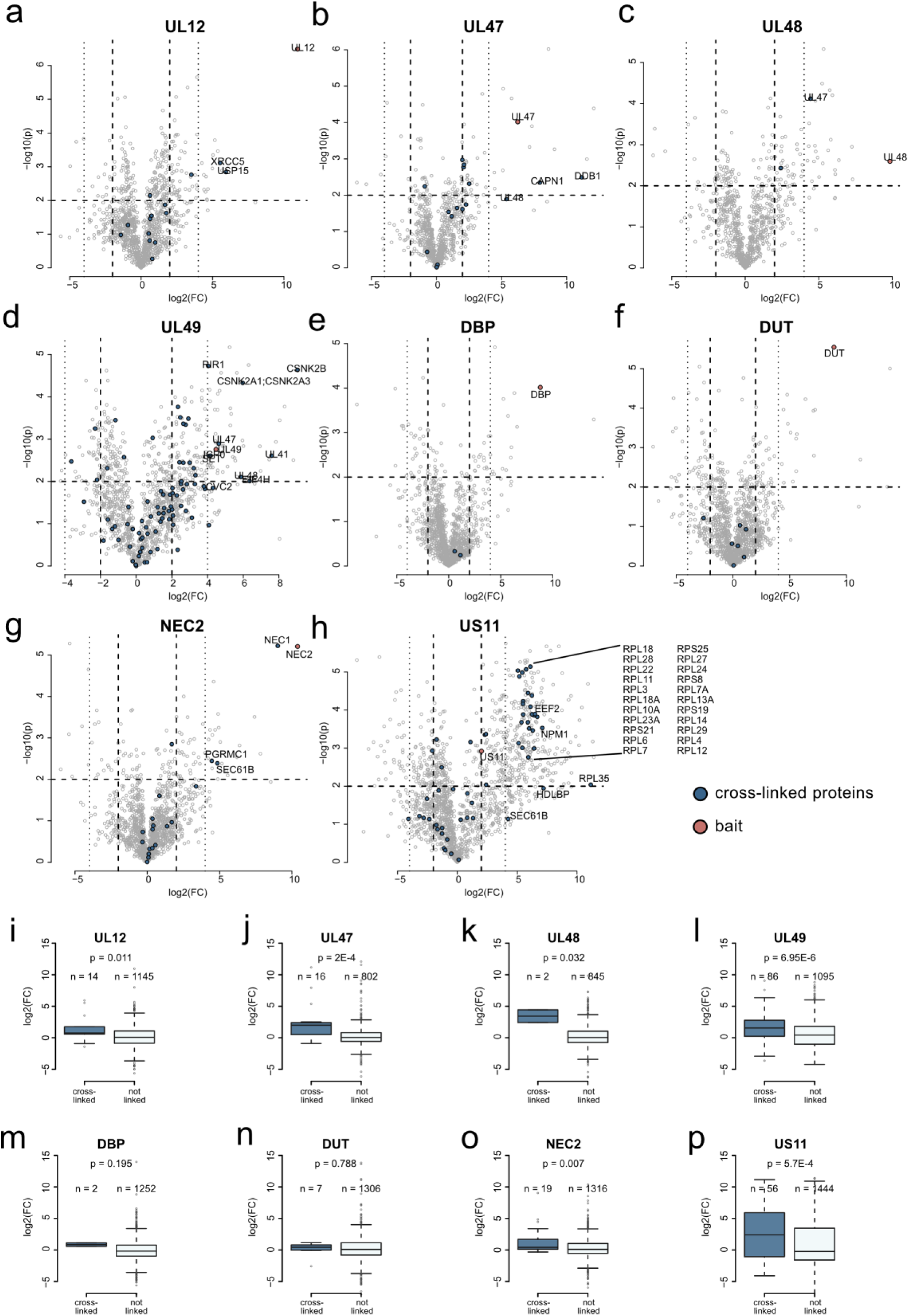
Volcano plot analysis of AP-MS experiments. **(a-h)** Each volcano plot gives the log2 fold-change as an estimate of the effect size (x-axis) and -log10 P-value (y-axis) as an estimate for the significance for protein interactions to a specific bait. P-values are based on two-sided t-tests without multiple hypothesis correction and n=3 biological replicates. Proteins cross-linked to the respective bait when co-enriched with a log2 fold-change > 4 are highlighted and labeled with their Gene names. **(i-p)** Average co-enrichment levels of cross-linked or non-cross-linked proteins to the bait for the individual AP-MS experiments. P-values are based on two-sided wilcoxon rank sum tests.

**Extended Data Figure 5.**
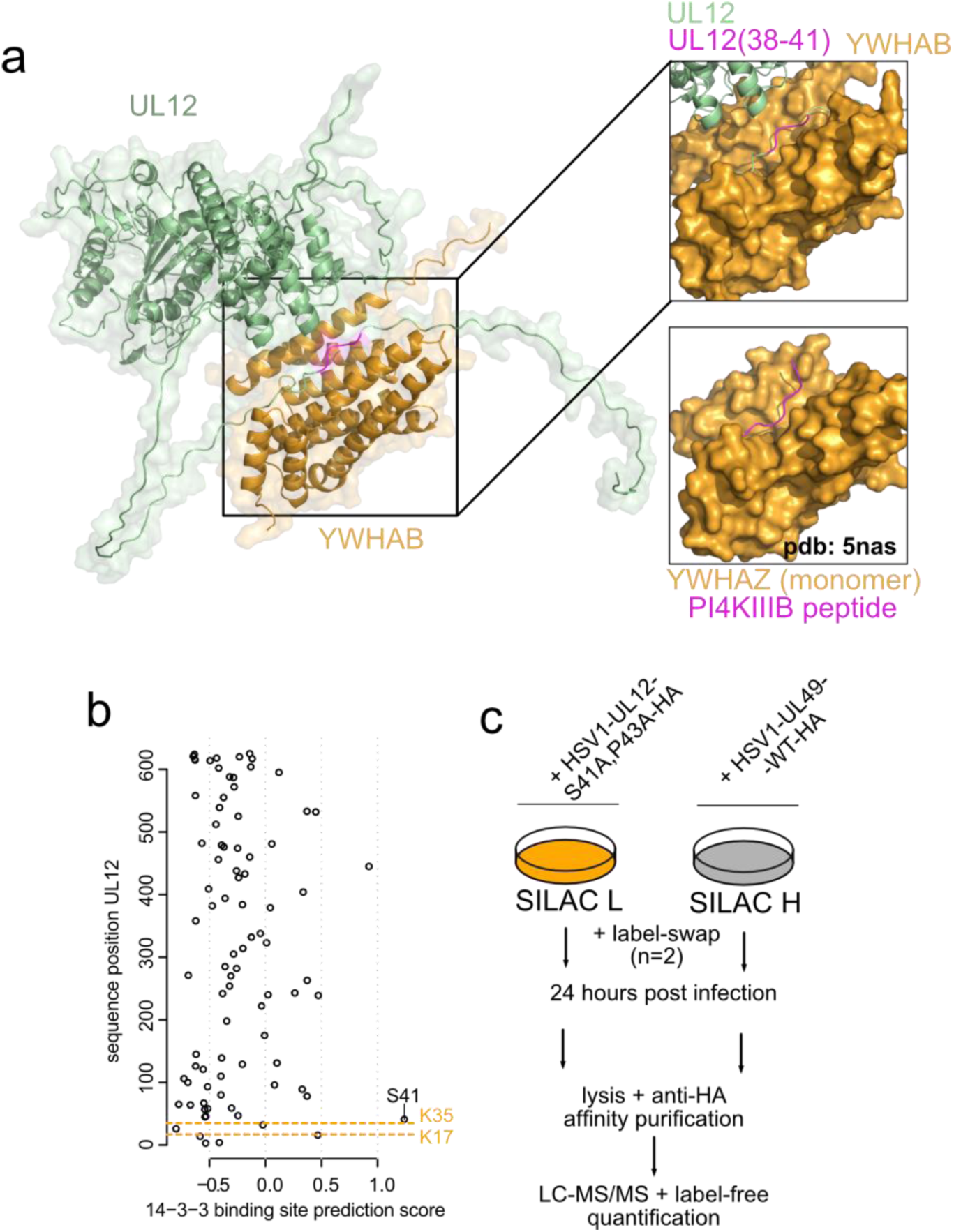
Interaction of 14-3-3 with UL12. **(a)** Predicted UL12-YWHAB dimer with UL12 (38-41) highlighted magenta, along with a comparison to crystallized structure of YHWHAZ and a 14-3-3 binding peptide of the PI4KIIIB protein. **(b)** Prediction of 14-3-3 binding site consensus score along the UL12 primary sequence based on 14-3-3pred ^68^. The best-scoring site S41 and the lysines cross-linked to 14-3-3 are highlighted. **(c)** Experimental design for SILAC-based comparative AP-MS of widltype-UL12 to S41A, P43A-UL12.

**Extended Data Figure 6.**
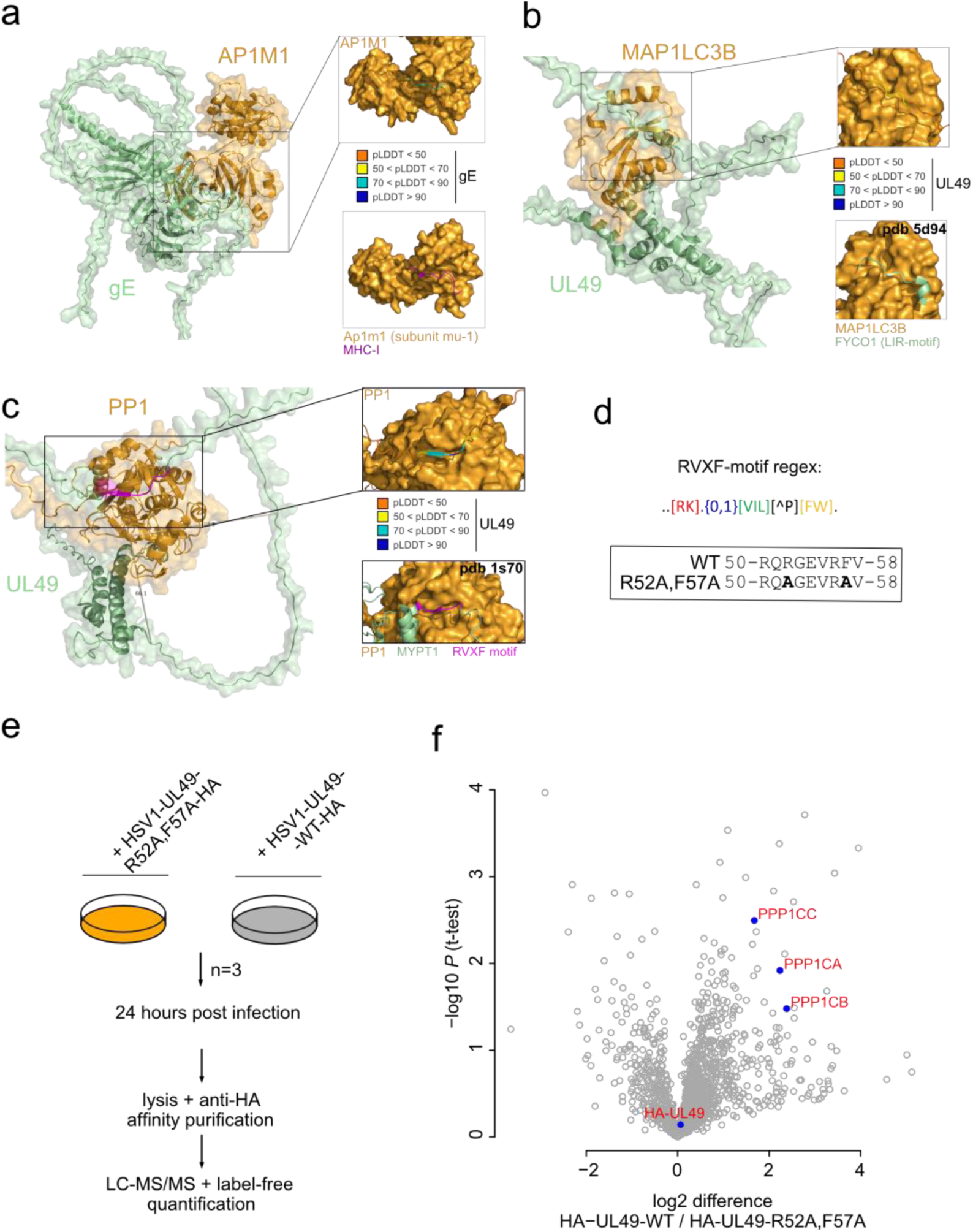
Interactions mediated through IDRs. **(a-c)** Examples of interactions putatively mediated through IDRs of gE-AP1M1 (a), UL49-MAP1LC3B (b) and UL49-PPP1CA (c). Insets show confidence of AF2 prediction based on pLDDT. Comparisons to experimentally solved structures are shown below the insets. **(d)** Regular expression for RVXF-type interaction motifs to PP-1. Boxed is the amino acid sequence of UL49 having a non-canonical RVXF-motif. Mutation of the non-canonical RVXF-motif in UL49 creates R52A, F57A-UL49. **(e)** Experimental design for label-free AP-MS experiments comparing HSV1-HA-UL49-WT to HSV1-HA-UL49-R52A, F57A. Experiment was performed in n=3 biological replicates. **(f)** AP-MS experiments directly comparing the interactome of WT UL49 to mutant UL49 in a label-free set-up based on n=3 biological replicates from infected cells. The bait (HA-UL49) and the three different catalytic subunits of PP-1, which can all interact with RVXF-type motifs, are labeled.

